## Supplementary material for "Conformations of a highly expressed Z19 α-zein studied with AlphaFold2 and MD simulations": S1 Appendix

### S1 Appendix. Overview of major α-zein sequences

An overview of major α-zein sequences and their names was given in the 1985 study by Marks *et al.* [1] on zein cDNA clones from the endosperm of the W64A maize inbred line. They used cross-hybridization to identify 3 cDNA clone groups (cZ22A, cZ22B, and cZ22C) for Z22 zeins and 5 cDNA clone groups (cZ19A, cZ19B, cZ19AB, cZ19C, cZ19D) for Z19 zeins. The authors sequenced at least one random cDNA clone from each group and named these sequences by appending integers to their group name (e.g., cZ19C1, cZ19D2, etc). These names are still used, but alternative names from both later and previous studies coexist and can give rise to confusion [2]. By presenting these sequences alongside previously published ones (gZ19AB1, cZ22A1 formerly known as pZ22.1, and cZ22B1 formerly known as pZ22.3) the article effectively summarized key α-zein sequences.

More recently, genomic analysis of endosperm in the B73 maize inbred line also found only a small number of expressed α-zein genes [2]. Importantly, the previous sequences [1] and new related sequences all fell into three subfamilies, designated "19 kD α-zein B," "19 kD α-zein D," and "22 kD α-zein," respectively. Sequences from different subfamilies shared 56% to 67% nucleotide identity (40% to 55% amino acid identity), allowing their differentiation through DNA/RNA-probe hybridization. However, sequences within the same subfamily had a higher nucleotide identity of 80% to 99% (75% to 95% amino acid identity), rendering them indistinguishable using this method. The three most highly expressed randomly chosen α-zein clones were αz19b1 (~16%), αz19b3 (=cZ19C2) (~6%), and αz22z1 (~5%), where percentages denote expressed sequence tags. The α-zein sequences from the above studies are listed in Table S1, which also includes the related Z19 zein M6 reported by Viotti in 1985 [3]. The table shows the amino acid composition in percent of the 17 zein sequences (Table S2 is the corresponding table with amino acid counts instead of percentages).

Theses tables show that all sequences lack Lys, and Trp is found only in αz22z4 and cZ22C2 in small amounts (0.4%). Z22 zeins have more Met (1.2 % to 2.0 %) relative to Z19 zeins (0.0 % to 0.5 %) which allows discrimination between the two groups, as does the lower Phe content of Z22 (3.3 % to 3.7 %) relative to Z19 (5.0 % to 6.3 %). Similarly, the 19 kD D-subfamily (αz19D2, cZ19D1, and αz19D1) can be identified by lower Ser (3.2 % - 4.1 %) compared to the remaining zeins (6.1 % - 8.2 %). PCA analysis of the sequences each represented by their content of each amino acids in percent (i.e., disregarding explicit sequence information), showed clustering into the three subfamilies, see Fig A.

**Table A.** Amino acid composition (percentage, colored by value) of α-zein sequences excluding signal peptides from [1-3].

| Name | Subfamily | UniProt | Length | MW | Cluster | ALA | ARG | ASN | ASP | CYS | GLN | GLU | GLY | HIS | ILE | LEU | MET | PHE | PRO | SER | THR | TRP | TYR | VAL |
|---|---|---|---|---|---|---|---|---|---|---|---|---|---|---|---|---|---|---|---|---|---|---|---|---|
| αz22z4 | 22-kD | Q946V4 | 246 | 26923 | 0 (100.00 % | 14.6 | 0.8 | 4.9 | 0.0 | 0.4 | 21.5 | 0.8 | 1.6 | 1.2 | 4.9 | 17.9 | 1.6 | 3.3 | 8.5 | 6.1 | 2.8 | 0.4 | 3.3 | 5.3 |
| cZ22B1 | 22-kD | P04698 | 246 | 26950 | 0 (90.24 %) | 13.8 | 1.6 | 5.3 | 0.0 | 0.4 | 20.7 | 0.8 | 0.8 | 1.2 | 4.5 | 17.1 | 2.0 | 3.3 | 8.9 | 6.9 | 2.8 | 0.0 | 2.8 | 6.9 |
| αz22z3 | 22-kD | O48966 | 245 | 26753 | 0 (91.02 %) | 13.9 | 1.2 | 5.3 | 0.0 | 0.4 | 20.8 | 0.4 | 1.2 | 1.2 | 4.9 | 17.1 | 1.6 | 3.7 | 9.0 | 6.9 | 3.3 | 0.0 | 2.9 | 6.1 |
| cZ22C2 | 22-kD | P06679 | 245 | 26847 | 0 (92.65 %) | 12.7 | 0.8 | 4.9 | 0.4 | 0.4 | 21.6 | 0.8 | 2.0 | 0.8 | 4.1 | 17.6 | 1.2 | 3.7 | 9.0 | 6.9 | 2.9 | 0.4 | 3.3 | 6.5 |
| cZ22A1/αz22z1 | 22-kD | P04700 | 242 | 26360 | 0 (91.32 %) | 14.0 | 0.8 | 5.0 | 0.0 | 0.4 | 20.7 | 0.4 | 1.7 | 1.2 | 3.3 | 18.2 | 1.7 | 3.7 | 9.1 | 6.6 | 3.3 | 0.0 | 3.3 | 6.6 |
| gZ19AB1 | 19-kD B | P04704 | 214 | 23498 | 1 (100.00 %) | 14.5 | 1.4 | 4.2 | 0.5 | 0.9 | 18.2 | 0.5 | 1.9 | 1.4 | 4.2 | 20.6 | 0.5 | 6.1 | 9.8 | 7.0 | 1.9 | 0.0 | 3.7 | 2.8 |
| αz19B1 | 19-kD B | Q946V6 | 213 | 23358 | 1 (92.02 %) | 12.7 | 0.9 | 4.7 | 0.0 | 0.9 | 19.2 | 0.5 | 2.3 | 0.9 | 4.2 | 19.7 | 0.0 | 6.1 | 10.8 | 7.5 | 3.3 | 0.0 | 3.8 | 2.3 |
| cZ19B1 | 19-kD B | P06675 | 213 | 23358 | 1 (91.08%) | 11.7 | 0.9 | 4.7 | 0.0 | 0.9 | 19.2 | 0.5 | 2.8 | 0.9 | 4.2 | 19.7 | 0.0 | 6.1 | 10.8 | 8.0 | 2.3 | 0.0 | 3.8 | 3.3 |
| cZ19A2 | 19-kD B | P06674 | 212 | 23192 | 1 (92.45%) | 14.2 | 0.9 | 4.2 | 0.9 | 0.9 | 18.4 | 0.5 | 2.4 | 1.4 | 4.2 | 19.3 | 0.0 | 6.1 | 10.8 | 6.1 | 2.8 | 0.0 | 3.8 | 2.8 |
| cZ19C2/αz19B3 | 19-kD B | P06677 | 219 | 24088 | 2 (100.00%) | 13.7 | 1.4 | 4.6 | 0.5 | 0.9 | 19.6 | 0.5 | 0.9 | 0.5 | 5.5 | 19.6 | 0.0 | 5.5 | 10.0 | 7.8 | 3.2 | 0.0 | 3.7 | 2.3 |
| cZ19C1 | 19-kD B | P06676 | 219 | 24092 | 2 (99.09 %) | 13.7 | 1.4 | 4.6 | 0.5 | 0.9 | 19.6 | 0.5 | 0.9 | 0.5 | 4.6 | 19.6 | 0.5 | 5.5 | 10.0 | 7.8 | 3.2 | 0.0 | 3.7 | 2.7 |
| zeinM6 | 19-kD B | P04702 | 219 | 24060 | 2 (97.72%) | 13.2 | 1.4 | 4.6 | 0.5 | 1.4 | 19.6 | 0.5 | 0.9 | 0.5 | 5.5 | 19.6 | 0.0 | 5.0 | 9.6 | 8.2 | 3.2 | 0.0 | 3.7 | 2.7 |
| αz19D2 | 19-kD D | Q946V7 | 220 | 24708 | 3 (100.00 %) | 12.3 | 2.3 | 5.9 | 0.5 | 0.5 | 21.4 | 0.9 | 1.8 | 0.5 | 5.5 | 17.7 | 0.5 | 5.9 | 8.2 | 3.2 | 3.6 | 0.0 | 4.1 | 5.5 |
| cZ19D1 | 19-kD D | P06678 | 219 | 24580 | 3 (97.26%) | 11.4 | 2.3 | 5.5 | 0.9 | 0.5 | 21.5 | 0.5 | 1.8 | 0.5 | 5.0 | 17.8 | 0.0 | 5.9 | 8.2 | 3.7 | 4.1 | 0.0 | 4.1 | 6.4 |
| αz19B2 | 19-kD B | Q548E7 | 246 | 27130 | 4 (100.00 %) | 14.2 | 1.6 | 4.5 | 0.4 | 0.8 | 19.1 | 0.4 | 1.6 | 2.0 | 4.1 | 21.1 | 0.4 | 5.3 | 8.9 | 6.5 | 1.6 | 0.0 | 3.7 | 3.7 |
| αz19D1 | 19-kD D | Q946V8 | 222 | 24820 | 5 (100.00 %) | 11.3 | 1.4 | 5.0 | 0.0 | 0.5 | 20.3 | 0.5 | 2.7 | 1.8 | 5.4 | 17.1 | 0.5 | 6.3 | 8.6 | 4.1 | 4.5 | 0.0 | 4.5 | 5.9 |
| αz22z5 | 22-kD | Q9SYT3 | 245 | 26710 | 6 (100 %) | 14.7 | 1.2 | 5.3 | 0.4 | 0.4 | 20.4 | 0.8 | 1.6 | 0.8 | 4.9 | 18.0 | 1.2 | 3.7 | 7.8 | 6.5 | 3.3 | 0.0 | 3.3 | 5.7 |

Name, zein name used in the original studies ('/' is used as a name separator when the two studies use different names to refer to the same sequence); Subfamily, subfamily names used in the study [2]; UniProtKB, UniProtKB ID for the sequence; Length, number of amino acids without signal peptide; MW, calculated molecular weight; Cluster, cluster number assigned by CD-HIT [4] for sequences with at least 90% identitiy followed by the in-cluster sequence identity (percentage) with the cluster representative in brackets.

**Table B.** Amino acid composition (number of amino acids, colored by value) of α-zein sequences from [1-3]. Signal peptides are excluded.

| Name | Subfamily | UniProt | Length | MW | Cluster | ALA | ARG | ASN | ASP | CYS | GLN | GLU | GLY | HIS | ILE | LEU | MET | PHE | PRO | SER | THR | TRP | TYR | VAL |
|---|---|---|---|---|---|---|---|---|---|---|---|---|---|---|---|---|---|---|---|---|---|---|---|---|
| αz22z4 | 22-kD | Q946V4 | 246 | 26923 | 0 (100.00 % | 36 | 2 | 12 | 0 | 1 | 53 | 2 | 4 | 3 | 12 | 44 | 4 | 8 | 21 | 15 | 7 | 1 | 8 | 13 |
| cZ22B1 | 22-kD | P04698 | 246 | 26950 | 0 (90.24 %) | 34 | 4 | 13 | 0 | 1 | 51 | 2 | 2 | 3 | 11 | 42 | 5 | 8 | 22 | 17 | 7 | 0 | 7 | 17 |
| αz22z3 | 22-kD | O48966 | 245 | 26753 | 0 (91.02 %) | 34 | 3 | 13 | 0 | 1 | 51 | 1 | 3 | 3 | 12 | 42 | 4 | 9 | 22 | 17 | 8 | 0 | 7 | 15 |
| cZ22C2 | 22-kD | P06679 | 245 | 26847 | 0 (92.65 %) | 31 | 2 | 12 | 1 | 1 | 53 | 2 | 5 | 2 | 10 | 43 | 3 | 9 | 22 | 17 | 7 | 1 | 8 | 16 |
| cZ22A1/αz22z1 | 22-kD | P04700 | 242 | 26360 | 0 (91.32 %) | 34 | 2 | 12 | 0 | 1 | 50 | 1 | 4 | 3 | 8 | 44 | 4 | 9 | 22 | 16 | 8 | 0 | 8 | 16 |
| gZ19AB1 | 19-kD B | P04704 | 214 | 23498 | 1 (100.00 %) | 31 | 3 | 9 | 1 | 2 | 39 | 1 | 4 | 3 | 9 | 44 | 1 | 13 | 21 | 15 | 4 | 0 | 8 | 6 |
| αz19B1 | 19-kD B | Q946V6 | 213 | 23358 | 1 (92.02 %) | 27 | 2 | 10 | 0 | 2 | 41 | 1 | 5 | 2 | 9 | 42 | 0 | 13 | 23 | 16 | 7 | 0 | 8 | 5 |
| cZ19B1 | 19-kD B | P06675 | 213 | 23358 | 1 (91.08%) | 25 | 2 | 10 | 0 | 2 | 41 | 1 | 6 | 2 | 9 | 42 | 0 | 13 | 23 | 17 | 5 | 0 | 8 | 7 |
| cZ19A2 | 19-kD B | P06674 | 212 | 23192 | 1 (92.45%) | 30 | 2 | 9 | 2 | 2 | 39 | 1 | 5 | 3 | 9 | 41 | 0 | 13 | 23 | 13 | 6 | 0 | 8 | 6 |
| cZ19C2/αz19B3 | 19-kD B | P06677 | 219 | 24088 | 2 (100.00%) | 30 | 3 | 10 | 1 | 2 | 43 | 1 | 2 | 1 | 12 | 43 | 0 | 12 | 22 | 17 | 7 | 0 | 8 | 5 |
| cZ19C1 | 19-kD B | P06676 | 219 | 24092 | 2 (99.09 %) | 30 | 3 | 10 | 1 | 2 | 43 | 1 | 2 | 1 | 10 | 43 | 1 | 12 | 22 | 17 | 7 | 0 | 8 | 6 |
| zeinM6 | 19-kD B | P04702 | 219 | 24060 | 2 (97.72%) | 29 | 3 | 10 | 1 | 3 | 43 | 1 | 2 | 1 | 12 | 43 | 0 | 11 | 21 | 18 | 7 | 0 | 8 | 6 |
| αz19D2 | 19-kD D | Q946V7 | 220 | 24708 | 3 (100.00 %) | 27 | 5 | 13 | 1 | 1 | 47 | 2 | 4 | 1 | 12 | 39 | 1 | 13 | 18 | 7 | 8 | 0 | 9 | 12 |
| cZ19D1 | 19-kD D | P06678 | 219 | 24580 | 3 (97.26%) | 25 | 5 | 12 | 2 | 1 | 47 | 1 | 4 | 1 | 11 | 39 | 0 | 13 | 18 | 8 | 9 | 0 | 9 | 14 |
| αz19B2 | 19-kD B | Q548E7 | 246 | 27130 | 4 (100.00 %) | 35 | 4 | 11 | 1 | 2 | 47 | 1 | 4 | 5 | 10 | 52 | 1 | 13 | 22 | 16 | 4 | 0 | 9 | 9 |
| αz19D1 | 19-kD D | Q946V8 | 222 | 24820 | 5 (100.00 %) | 25 | 3 | 11 | 0 | 1 | 45 | 1 | 6 | 4 | 12 | 38 | 1 | 14 | 19 | 9 | 10 | 0 | 10 | 13 |
| αz22z5 | 22-kD | Q9SYT3 | 245 | 26710 | 6 (100 %) | 36 | 3 | 13 | 1 | 1 | 50 | 2 | 4 | 2 | 12 | 44 | 3 | 9 | 19 | 16 | 8 | 0 | 8 | 14 |

Name, zein name used in the original studies ('/' is used as a name separator when the two studies use different names to refer to the same sequence); Subfamily, subfamily names used in the study [2]; UniProtKB, UniProtKB ID for the sequence; Length, number of amino acids without signal peptide; MW, calculated molecular weight; Cluster, cluster number assigned by CD-HIT [4] for sequences with at least 90% identitiy followed by the in-cluster sequence identity (percentage) with the cluster representative in brackets.

**Table C. α-zein sequences from previous studies of maize endosperm**

| Name/Alt. name | UniProtKB name(s) | UniProtKB ID | Study, Maize line | Evidence level |
|---|---|---|---|---|
| ZeinA30 | 19 kDa zein A30 | P02859 | [5], IHP | Transcript |
| ZeinA20 | 19 kDa zein A20 | P04703 | [6], IHP | Transcript |
| cZ22A1/αz22z1 | 22 kDa alpha-zein 16 | P04700 | [1], W64A/[2], B73 | Transcript |
| cZ22B1 | 22 kDa alpha-zein 4, Zein- alpha PZ22.3 | P04698 | [1], W64A | Transcript |
| cZ22C2 | 22 kDa alpha-zein 8b, Zein- alpha 22C2 | P06679 | [1], W64A | Transcript |
| gZ19AB1 | Zein- alpha ZG99, 19 kDa zein ZG99 | P04704 | [7], [1], W64A | Transcript |
| cZ19A2 | Zein-alpha 19A2, 19 kDa zein 19A2 | P06674 | [1], W64A | Transcript |
| cZ19B1 | Zein-alpha 19B1, 19 kDa zein 19B1 | P06675 | [1], W64A | Transcript |
| αz19B1 | 19kD alpha zein B1 | Q946V6 | [2], B73 | Transcript |
| αz19B2 | 19kD alpha zein B2 | Q548E7 | [2], B73 | Transcript |
| cZ19D1 | Zein-alpha 19D1, 19 kDa zein 19D1 | P06678 | [1], W64A | Transcript |
| cZ19C1 | Zein-alpha 19C1, 19 kDa zein 19C1 | P06676 | [1], W64A | Transcript |
| cZ19C2/αz19B3 | 19 kDa alpha-zein 19C2, Zein-alpha 19C2 | P06677/Q548E6 | [1], W64A/[2], B73 | **Protein** |
| ZeinM6 | 19 kDa zein M6 | P04702/ X02450 | [3], W64A | Transcript |
| αz19D1 | 19kD alpha zein D1 | Q946V8 | [2], B73 | Transcript |
| αz19D2 | 19kD alpha zein D2 | Q946V7 | [2], B73 | Transcript |
| αz22z3 | 22 kDa alpha-zein 4, PZ22.3 | O48966 | [2], B73 | **Protein** |
| αz22z4 | 22kD alpha zein 4 | Q946V4 | [2], B73 | Transcript |
| αz22z5 | 22kD alpha zein 5 | Q9SYT3 | [2], B73 | Transcript |

The relatedness of the protein sequences in Table A is illustrated by the tree in Fig 1, made with Mega11 [8]. The two main branches separate Z19 and Z22 zeins, while Z19 zeins branch further into the 3-member "19 kD D" subfamily and the "19 kD B" subfamily containing the 8 remaining Z19 zeins. A similar grouping was obtained by clustering the amino acid sequences with 90% identity using CD-HIT [4], giving 7 clusters indicated by indices 0 to 6 under "Cluster" in Table A.

The high homology of certain sequences when compared across independent studies, indicates that these studies may have converged to some extent in mapping highly expressed α-zeins from common maize variants. For instance, the mature protein sequence A30 (UniProtKB ID: P02859) differs by 6 amino acid changes from cz19B1 (UniProtKB ID: P06675), while A20 (UniProtKB ID: P04703) differs by 3 amino acid changes from cZ19C2 (UniProtKB ID: P06677).

### PCA on α-zein amino acid profiles

To examine if the subfamily grouping of zeins from hybridization experiments [1] could be approximated without explicit sequence information, a principal components analysis (PCA) was conducted where the zeins were represented only by their amino acid percentages (Table A). The scores plot (Fig A, left) shows three distinct clusters corresponding to the subfamilies 19-kD B, 19-kD D, and 22-kD from [9], with cZ19C2 studied here located centrally in the 19-kD B cluster.

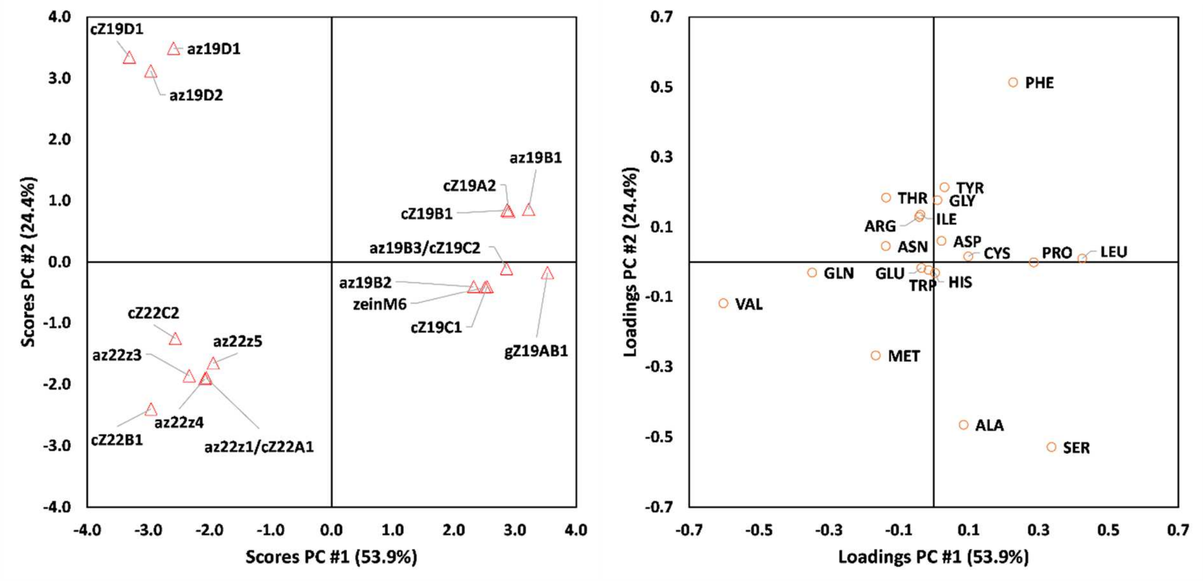


**Fig A. Principal component analysis for α-zeins in Table A represented by amino acid content in percent.** Scores (left) and loadings (right) for the two first principal components (PCs). Percent variance explained is given in the axis labels.

### References


1. Marks MD, Lindell JS, Larkins BA. Nucleotide sequence analysis of zein mRNAs from maize endosperm. Journal of Biological Chemistry. 1985;260(30):16451-9. doi: 10.1016/S0021-9258(17)36258-0.
2. Woo Y-M, Hu DW-N, Larkins BA, Jung R. Genomics Analysis of Genes Expressed in Maize Endosperm Identifies Novel Seed Proteins and Clarifies Patterns of Zein Gene Expression. The Plant Cell. 2001;13(10):2297-317. doi: 10.1105/tpc.010240.
3. Viotti A, Cairo G, Vitale A, Sala E. Each zein gene class can produce polypeptides of different sizes. The EMBO Journal. 1985;4(5):1103-10. doi: https://doi.org/10.1002/j.1460-2075.1985.tb03746.x.
4. Fu L, Niu B, Zhu Z, Wu S, Li W. CD-HIT: accelerated for clustering the next-generation sequencing data. Bioinformatics. 2012;28(23):3150-2. Epub 2012/10/13. doi: 10.1093/bioinformatics/bts565. PubMed PMID: 23060610; PubMed Central PMCID: PMCPMC3516142.
5. Geraghty D, Peifer MA, Rubenstein I, Messing J. The primary structure of a plant storage protein: zein. Nucleic Acids Research. 1981;9(19):5163-74. doi: 10.1093/nar/9.19.5163.
6. Geraghty DE, Messing J, Rubenstein I. Sequence analysis and comparison of cDNAs of the zein multigene family. Embo j. 1982;1(11):1329-35. Epub 1982/01/01. doi: 10.1002/j.1460-2075.1982.tb01318.x. PubMed PMID: 6897917; PubMed Central PMCID: PMCPMC553212.

7. Pedersen K, Devereux J, Wilson DR, Sheldon E, Larkins BA. Cloning and sequence analysis reveal structural variation among related zein genes in maize. Cell. 1982;29(3):1015-26. doi: https://doi.org/10.1016/0092-8674(82)90465-2.
8. Tamura K, Stecher G, Kumar S. MEGA11: Molecular Evolutionary Genetics Analysis Version 11. Molecular Biology and Evolution. 2021;38(7):3022-7. doi: 10.1093/molbev/msab120.
9. Marks MD, Larkins BA. Analysis of sequence microheterogeneity among zein messenger RNAs. Journal of Biological Chemistry. 1982;257(17):9976-83. doi: https://doi.org/10.1016/S0021-9258(18)33973-5.
