## Supplementary material for "Conformations of a highly expressed Z19 α-zein studied with AlphaFold2 and MD simulations": S1 Fig

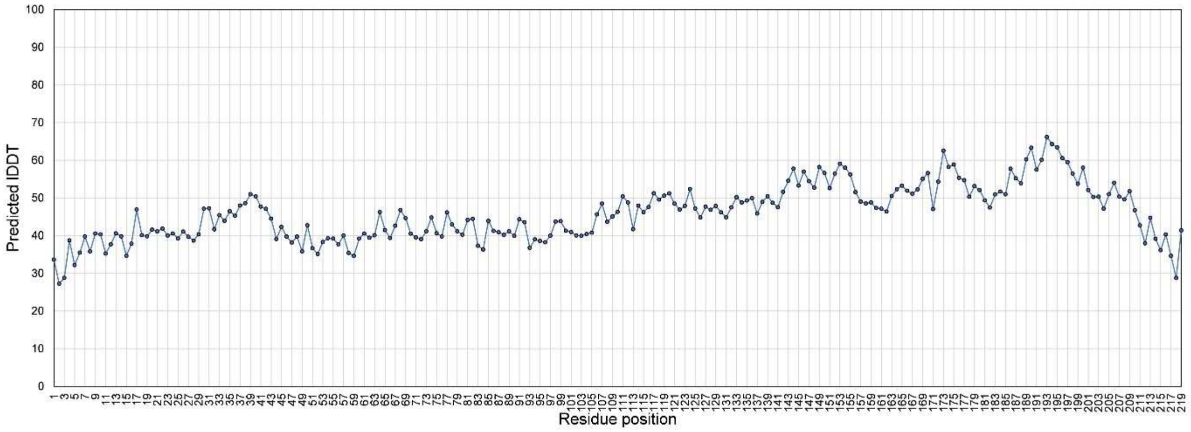


**AlphaFold2 per-residue estimate of confidence (plDDT) for the rank 1 model of α-zein cZ19C2 (UniProtKB ID: P06677).** The scale runs from 0 (minimum confidence) to 100 (maximum confidence).
