## Supplementary material for "Conformations of a highly expressed Z19 α-zein studied with AlphaFold2 and MD simulations": S2 Appendix

### S2 Appendix. Homology and *ab initio* models

To illustrate the uncertainty in α-zein homology models due to lack of templates, here homology modelling with I-TASSER [1] was carried out for the mature protein sequence of P06677 giving the dissimilar structures in Fig A, demonstrating the lack of convergence to a single prediction.

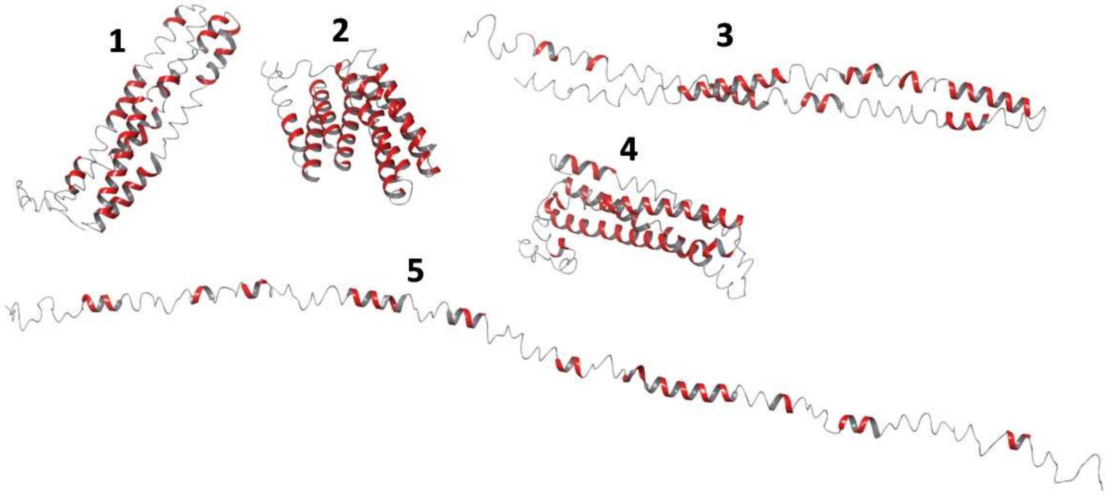


**Fig A. Homology models obtained using I-TASSER using the mature protein sequence of UniProt ID P06677.** C-scores for models 1, 2, 3, 4, 5 are -3.30, -2.31, -2.65, -3.99 and -4.13, respectively.

*Ab initio* modeling is an alternative to homology modeling, obviating the need for templates by combining structural sampling (e.g. Monte Carlo simulations) and physics- or knowledge-based potentials [2]. A recent example is the deep-learning contact assisted method C-QUARK [3]. When applied to P06677 herein this method generated five relatively similar models, as shown in Fig B. While the similarity of these models suggests convergence of the method, validation of these results is challenged by the lack of experimental zein structures.

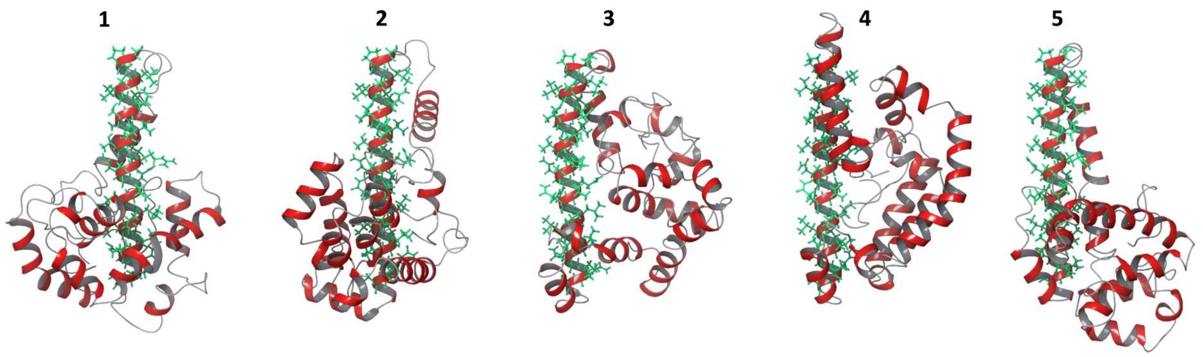


**Fig B.** ***Ab initio*** **models from C-QUARK using the mature protein sequence of UniProt ID P06677.** Residues S56 – A86 (green sticks) reside in a contiguous α-helix in all models. ProSA scores for models 1, 2, 3, 4, 5 are -6.35, -5.42, -6.09, -4.74, and -5.82, respectively.

The highest ranked models of P06677 from AlphaFold2, I-TASSER, and D-I-TASSER, C-QUARK are shown for comparison in Fig C. The models are clearly dissimilar, although the C-QUARK and AlphaFold2 models share that residues S56-A86 are α-helical.

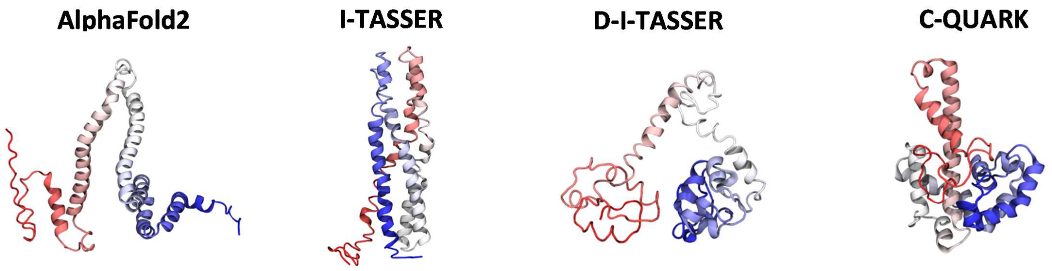


**Fig C. Models of the mature α-zein (UniProtKB ID: P06677) predicted with AlphaFold2, I-TASSER, D-I-TASSER, and C-QUARK.** Ribbon representations are colored by residue position (Red: N-terminal, Blue: C-terminal). Model quality estimates are given in Fig D.

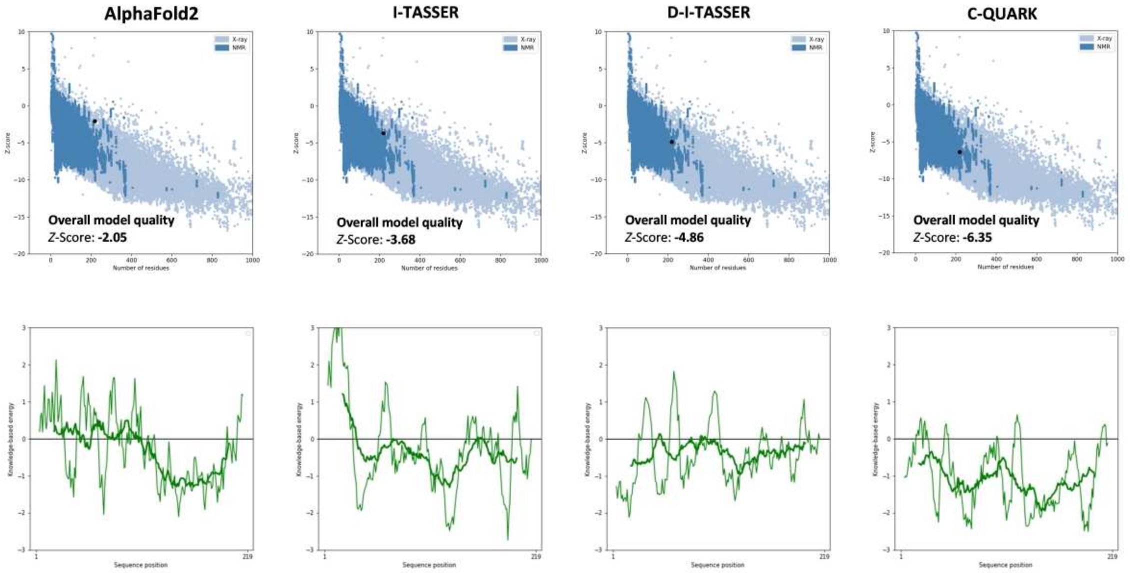


**Fig D. ProSA-web [4] quality checks for the AlphaFold2, I-TASSER, D-I-TASSER, and C-QUARK models of the mature α-zein in Fig C.**

### References


1. Yang J, Zhang Y. I-TASSER server: new development for protein structure and function predictions. Nucleic Acids Research. 2015;43(W1):W174-W81. doi: 10.1093/nar/gkv342.
2. Pearce R, Zhang Y. Toward the solution of the protein structure prediction problem. Journal of Biological Chemistry. 2021;297(1):100870. doi: https://doi.org/10.1016/j.jbc.2021.100870.
3. Mortuza SM, Zheng W, Zhang C, Li Y, Pearce R, Zhang Y. Improving fragment-based ab initio protein structure assembly using low-accuracy contact-map predictions. Nature Communications. 2021;12(1):5011. doi: 10.1038/s41467-021-25316-w.
4. Wiederstein M, Sippl MJ. ProSA-web: interactive web service for the recognition of errors in three-dimensional structures of proteins. Nucleic Acids Research. 2007;35(suppl_2):W407-W10. doi: 10.1093/nar/gkm290.
