## Supplementary material for "Conformations of a highly expressed Z19 α-zein studied with AlphaFold2 and MD simulations": S2 Fig

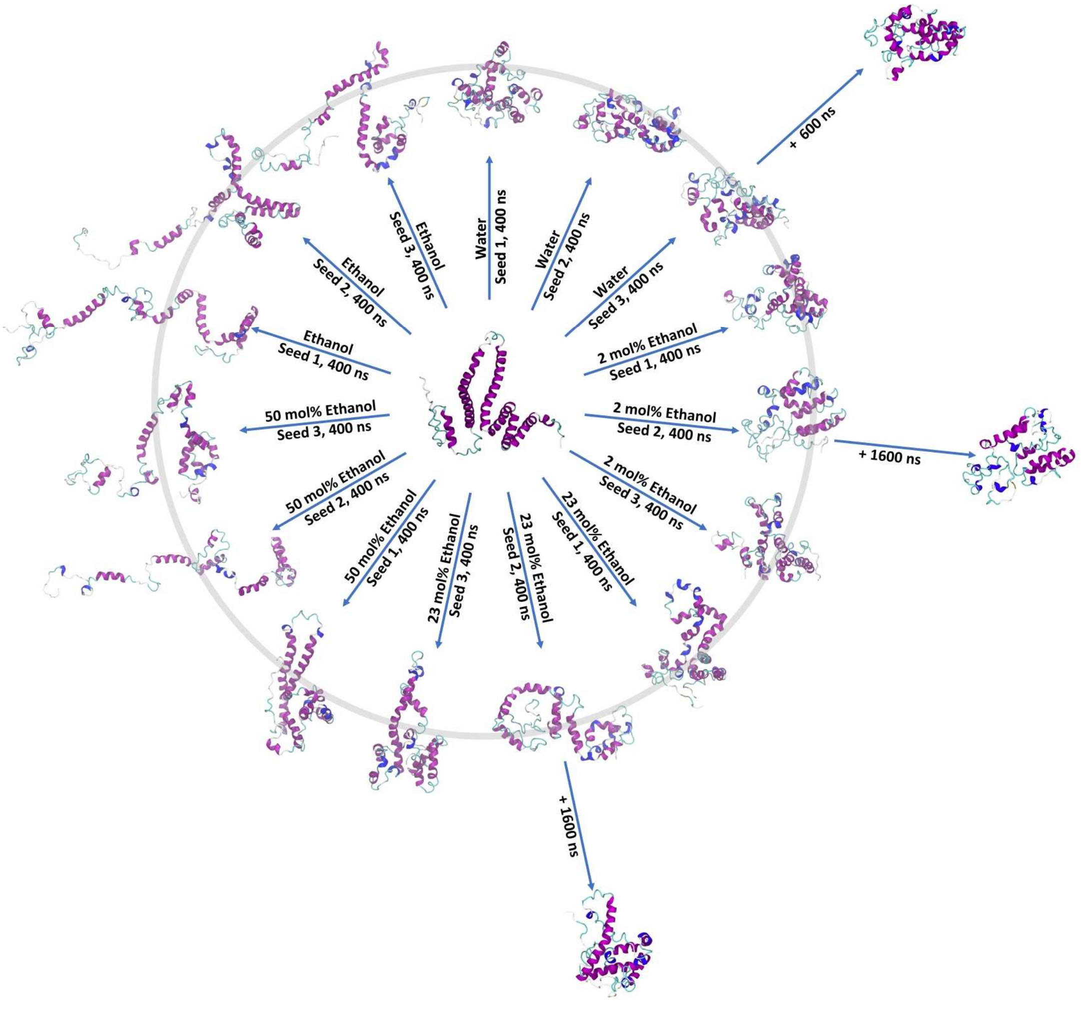


**Overview of the all-atom GROMACS simulations with the OPLS-AA/L + SPCE force fields.** 15 × 400 ns simulations of which 3 simulations were extended to at least 1 μs.
