## Supplementary material for "Conformations of a highly expressed Z19 α-zein studied with AlphaFold2 and MD simulations": S3 Appendix

### S3 Appendix. Additional simulation details

#### Table A. Protonation state of ionizable groups.

| Residue | Charge |
|---|---|
| N-terminal | +1 |
| CYS6 | 0 |
| CYS27 | 0 |
| GLU28 | -1 |
| ARG36 | +1 |
| ARG70 | +1 |
| ARG165 | +1 |
| ASP204 | -1 |
| HIS212 (Hε) | 0 |
| C-terminal | -1 |
| Total charge | +1 |

#### Solvent density in MD simulations

**Table B. Densities (kg/m$^3$) for 100 ps NPT equilibration with protein heavy atom position constraints in GROMACS.** Statistics over 50001 steps (501 points).

| System | Force field(s) | Average | Err.Est. | RMSD | Tot-Drift |
|---|---|---|---|---|---|
| **100 mol% water, Seed 1** | OPLS-AA/L; SPCE | 1002.51 | 0.22 | 1.93249 | 0.960957 |
| **100 mol% water, Seed 2** | OPLS-AA/L; SPCE | 1002.86 | 0.16 | 1.94758 | 0.901261 |
| **100 mol% water, Seed 3** | OPLS-AA/L; SPCE | 1002.76 | 0.13 | 1.85953 | 0.286725 |
| **2 mol% ethanol, Seed 1** | OPLS-AA/L; SPCE | 996.538 | 0.14 | 1.29564 | 0.74439 |
| **2 mol% ethanol, Seed 2** | OPLS-AA/L; SPCE | 996.371 | 0.13 | 1.23831 | 0.787757 |
| **2 mol% ethanol, Seed 3** | OPLS-AA/L; SPCE | 996.362 | 0.17 | 1.34724 | 0.575559 |
| **23 mol% ethanol, Seed 1** | OPLS-AA/L; SPCE | 933.376 | 0.69 | 2.60158 | 4.26358 |
| **23 mol% ethanol, Seed 2** | OPLS-AA/L; SPCE | 933.228 | 0.64 | 2.54067 | 4.20843 |
| **23 mol% ethanol, Seed 3** | OPLS-AA/L; SPCE | 933.098 | 0.58 | 2.43144 | 3.56655 |
| **50 mol% ethanol, Seed 1** | OPLS-AA/L; SPCE | 870.645 | 0.58 | 2.16417 | 3.39009 |
| **50 mol% ethanol, Seed 2** | OPLS-AA/L; SPCE | 870.482 | 0.65 | 2.16826 | 3.26837 |
| **50 mol% ethanol, Seed 3** | OPLS-AA/L; SPCE | 871.279 | 0.49 | 2.12289 | 2.5848 |
| **50 mol% ethanol, Seed 4** | OPLS-AA/L; SPCE | 870.905 | 0.48 | 2.0782 | 1.79602 |
| **100 mol% ethanol, Seed 1** | OPLS-AA/L; SPCE | 802.425 | 0.52 | 2.92551 | 3.18567 |

| | | | | | |
|---|---|---|---|---|---|
| **100 mol% ethanol, Seed 2** | OPLS-AA/L; SPCE | 802.631 | 0.63 | 2.49467 | 2.63548 |
| **100 mol% ethanol, Seed 3** | OPLS-AA/L; SPCE | 802.662 | 0.41 | 2.48046 | 1.78485 |
| **100 mol% water, Seed 1,** | ff99SB*-ILDN; TIP3P | 989.509 | 0.09 | 1.48081 | 0.242283 |
| **100 mol% water, Seed 2** | ff99SB*-ILDN; TIP3P | 989.438 | 0.13 | 1.4205 | 0.316504 |
| **24 mol% ethanol, Seed 1** | ff99SB*-ILDN; TIP3P | 900.301 | 0.15 | 1.54634 | 0.422864 |
| **24 mol% ethanol, Seed 2** | ff99SB*-ILDN; TIP3P | 900.897 | 0.17 | 1.65101 | -0.465899 |
