## Supplementary material for "Conformations of a highly expressed Z19 α-zein studied with AlphaFold2 and MD simulations": S3 Fig

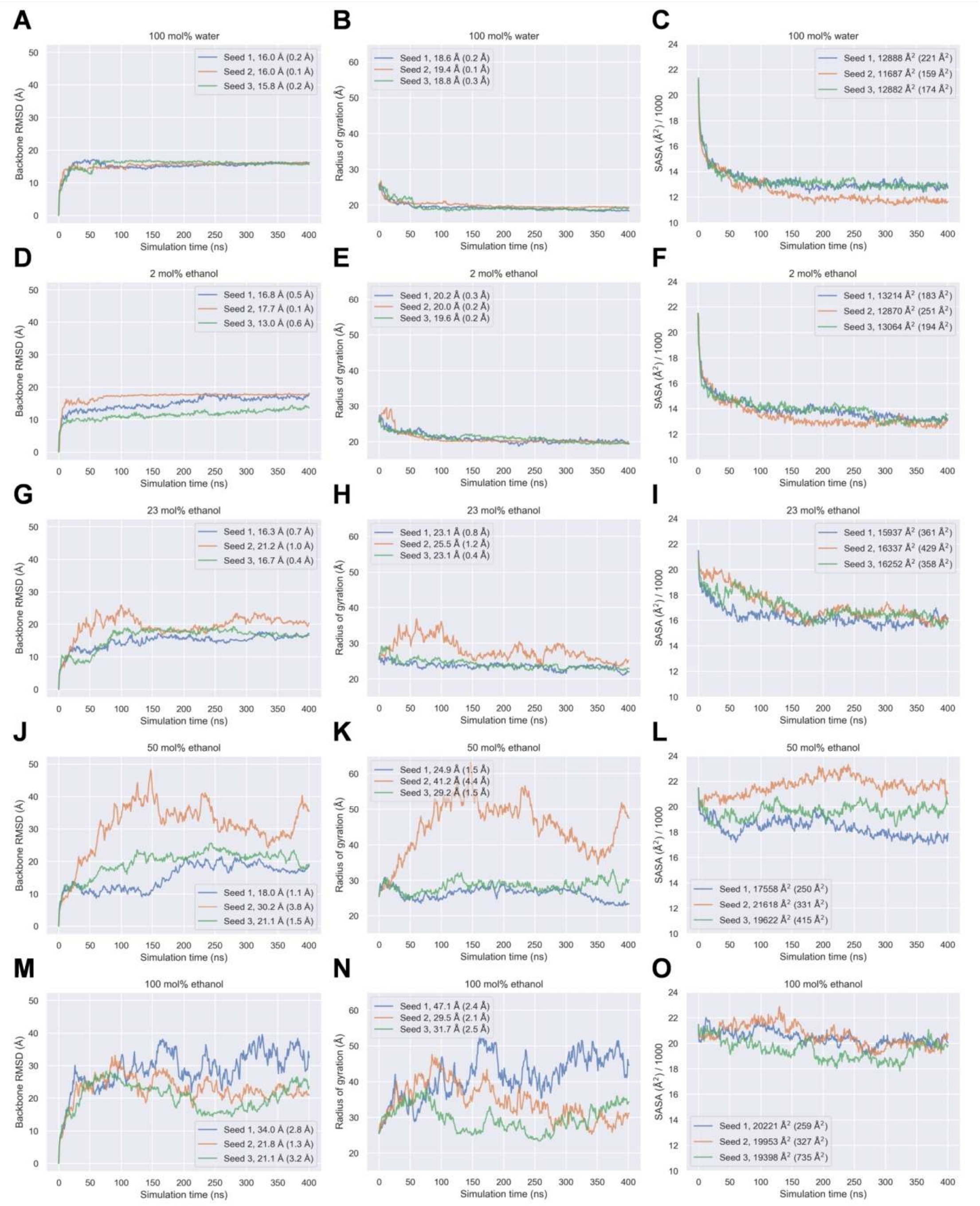


**Time series for 400 ns GROMACS all-atom MD simulations.** A, D, G, J, M: Backbone RMSD. B, E, H, K N: Radius of gyration. C, F, I, L O: SASA. Values given in the legend for each seed are averages over the last 100 ns of each simulation, with standard deviations given in brackets.
