## Supplementary material for "Conformations of a highly expressed Z19 α-zein studied with AlphaFold2 and MD simulations": S6 Fig

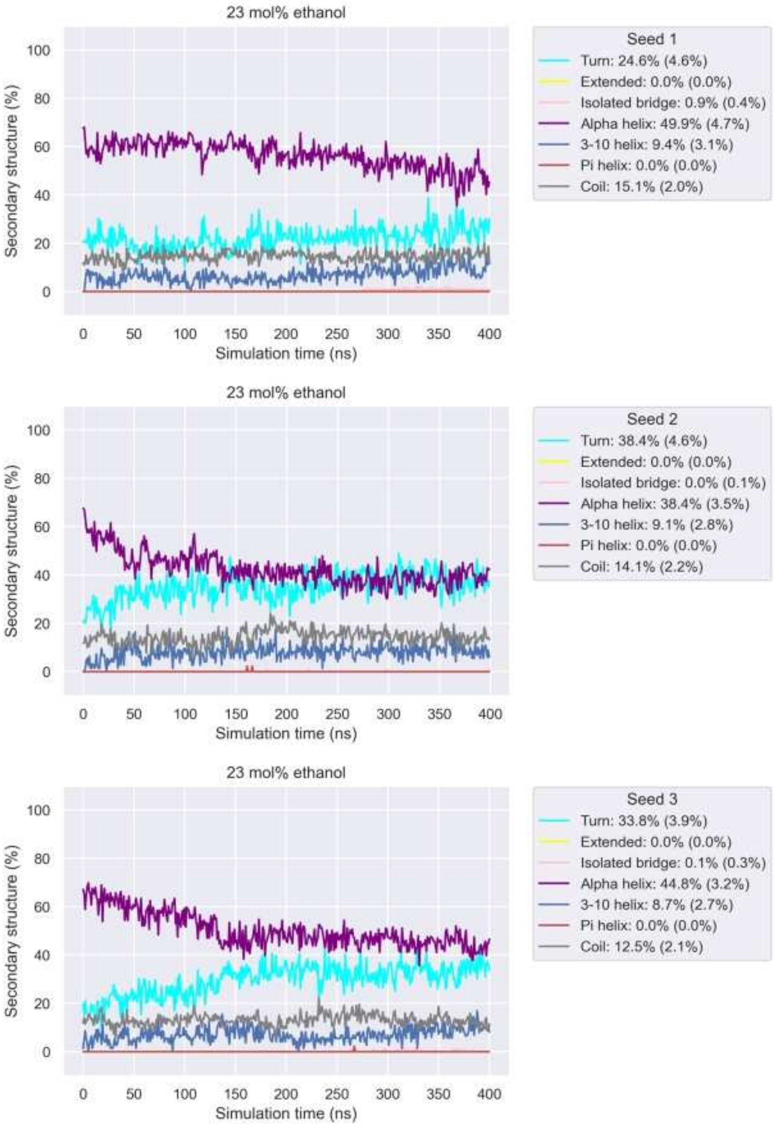


**Time series for secondary structure for 23 mol% ethanol 400 ns MD simulations.** Values given in the legend for each seed are averages over the last 100 ns of each simulation, with standard deviations given in brackets.
