## Supplementary material for "Conformations of a highly expressed Z19 α-zein studied with AlphaFold2 and MD simulations": S14 Fig

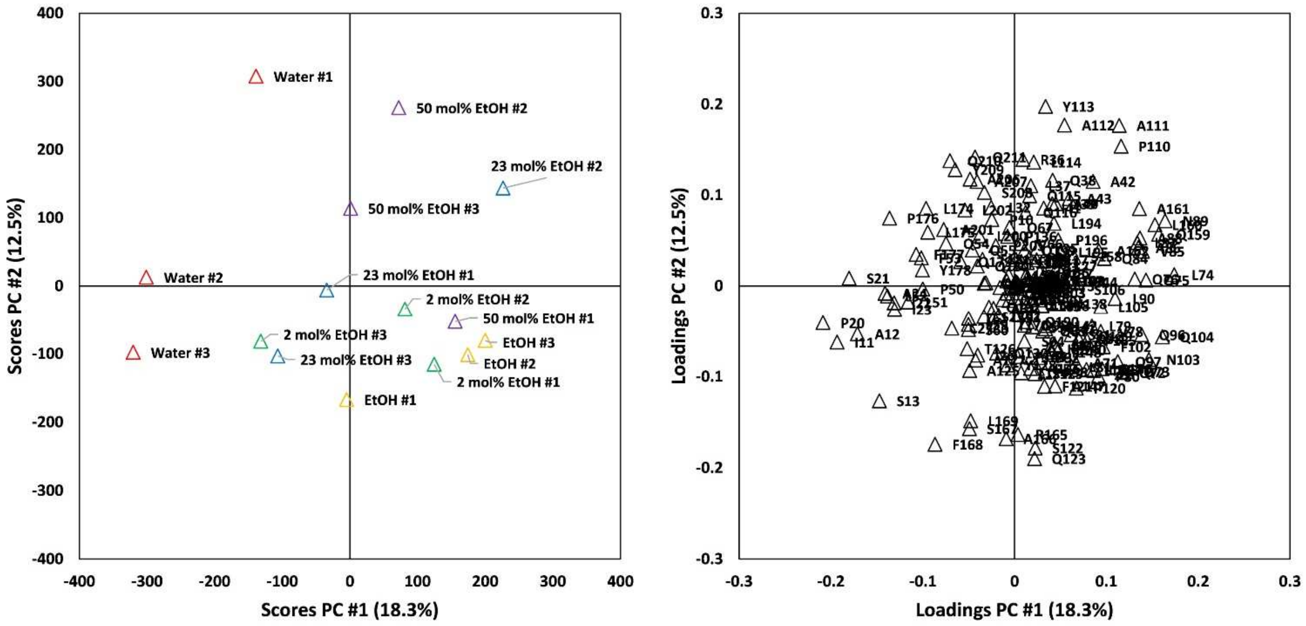


**Principal Component Analysis of Helicity Profiles from 400 ns MD simulations.** Left: Scores for the first two principal components. Right: Loadings for the two first principal components. The PCA was carried out on helicity profiles obtained from the last 100 ns of each 400 ns MD simulations by averaging the helicity per residue across the last 100 snapshots. Simulations are color coded based on ethanol concentration. The suffixes #1, #2, and #3 indicate different simulation replicates (seeds).
