## Supplementary material for "Conformations of a highly expressed Z19 α-zein studied with AlphaFold2 and MD simulations": S15 Fig

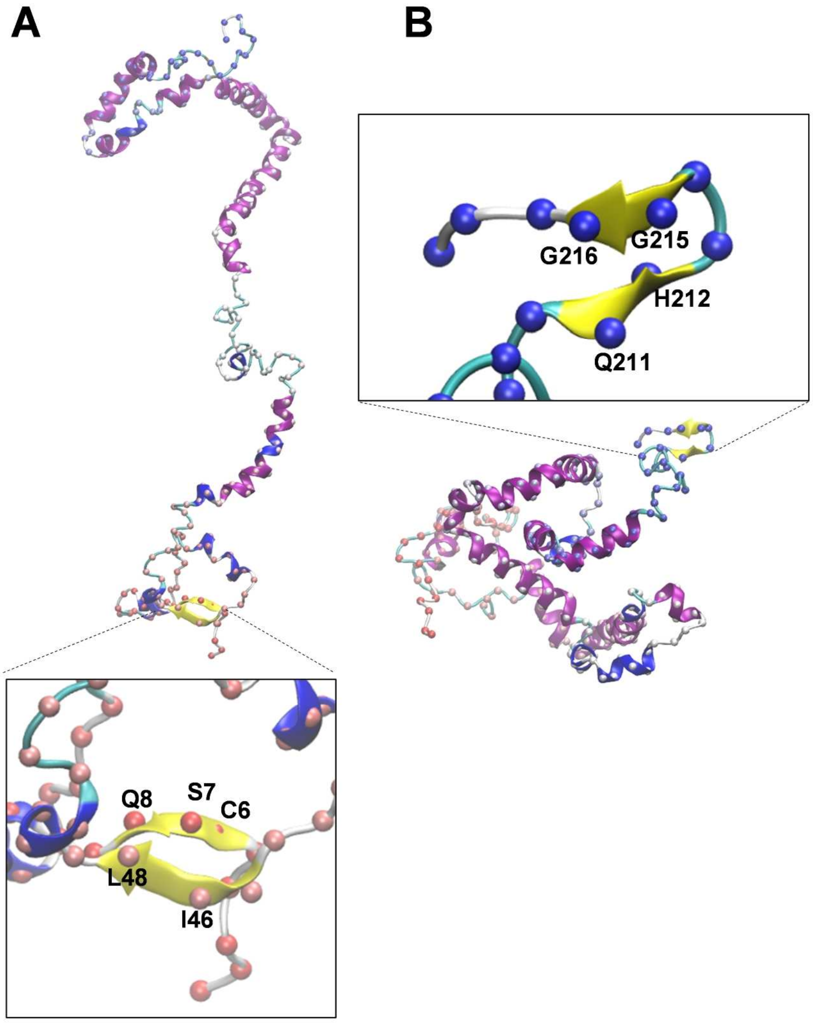


**Transient β-sheet formation in 400 ns ethanol simulations.** A: Seed 1, B: Seed 2. The snapshots are from $t$ = 346 ns in both cases.
