## Supplementary material for "Conformations of a highly expressed Z19 α-zein studied with AlphaFold2 and MD simulations": S16 Fig

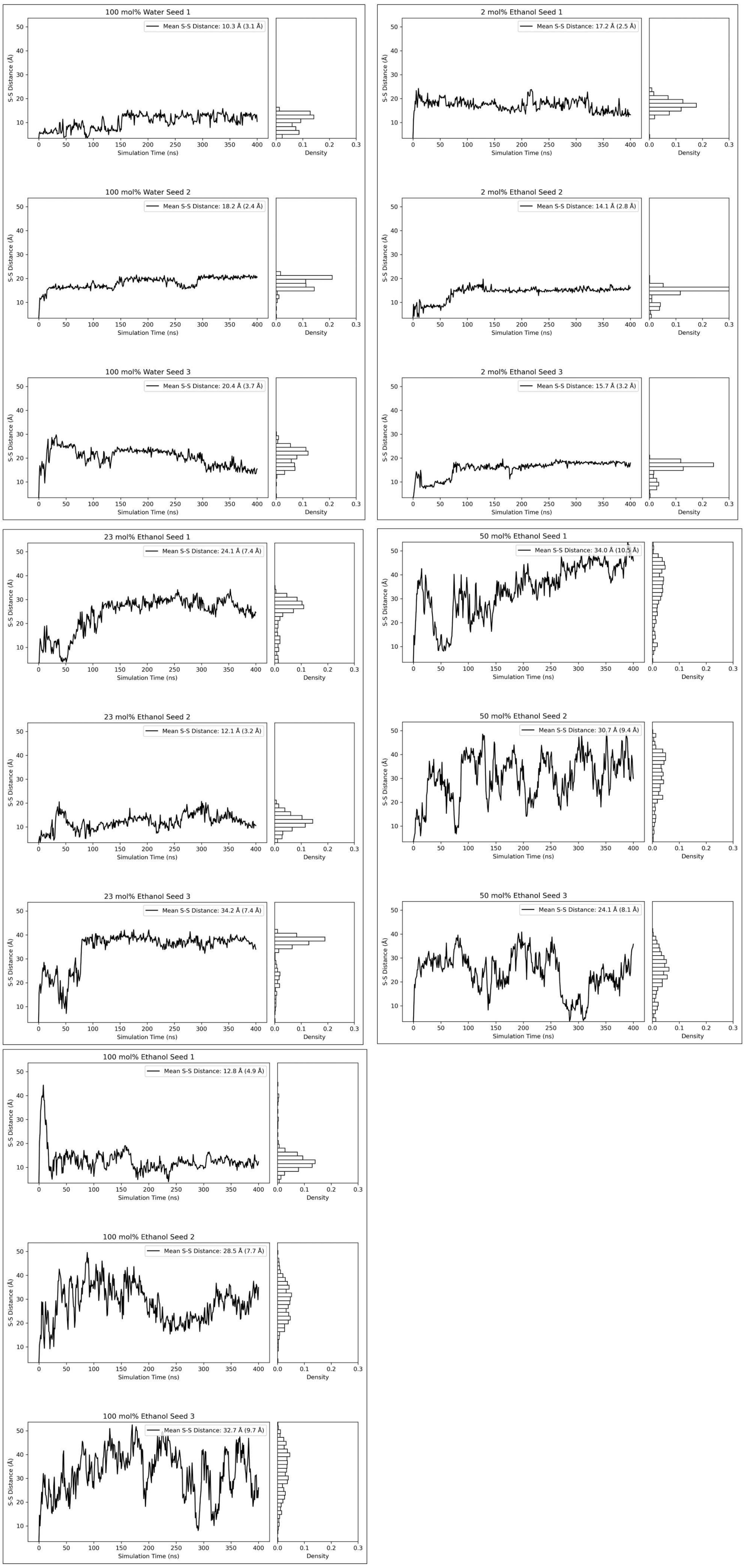
100 mol% Water Seed 1
Mean S-S Distance: 10.3 Å (3.1 Å)
100 mol% Water Seed 2
Mean S-S Distance: 18.2 Å (2.4 Å)
100 mol% Water Seed 3
Mean S-S Distance: 20.4 Å (3.7 Å)
2 mol% Ethanol Seed 1
Mean S-S Distance: 17.2 Å (2.5 Å)
2 mol% Ethanol Seed 2
Mean S-S Distance: 14.1 Å (2.8 Å)
2 mol% Ethanol Seed 3
Mean S-S Distance: 15.7 Å (3.2 Å)
23 mol% Ethanol Seed 1
Mean S-S Distance: 24.1 Å (7.4 Å)
23 mol% Ethanol Seed 2
Mean S-S Distance: 12.1 Å (3.2 Å)
23 mol% Ethanol Seed 3
Mean S-S Distance: 34.2 Å (7.4 Å)
50 mol% Ethanol Seed 1
Mean S-S Distance: 34.0 Å (10.5 Å)
50 mol% Ethanol Seed 2
Mean S-S Distance: 30.7 Å (9.4 Å)
50 mol% Ethanol Seed 3
Mean S-S Distance: 24.1 Å (8.1 Å)
100 mol% Ethanol Seed 1
Mean S-S Distance: 12.8 Å (4.9 Å)
100 mol% Ethanol Seed 2
Mean S-S Distance: 28.5 Å (7.7 Å)
100 mol% Ethanol Seed 3
Mean S-S Distance: 32.7 Å (9.7 Å)
S-S Distance (Å)
Simulation Time (ns)
Density

**Distance between sulfur atoms in the two cysteines across all 400 ns MD trajectories.** Mean and standard deviations in this plot are calculated using the entire 400 ns range.
