## Supplementary material for "Conformations of a highly expressed Z19 α-zein studied with AlphaFold2 and MD simulations": S17 Fig

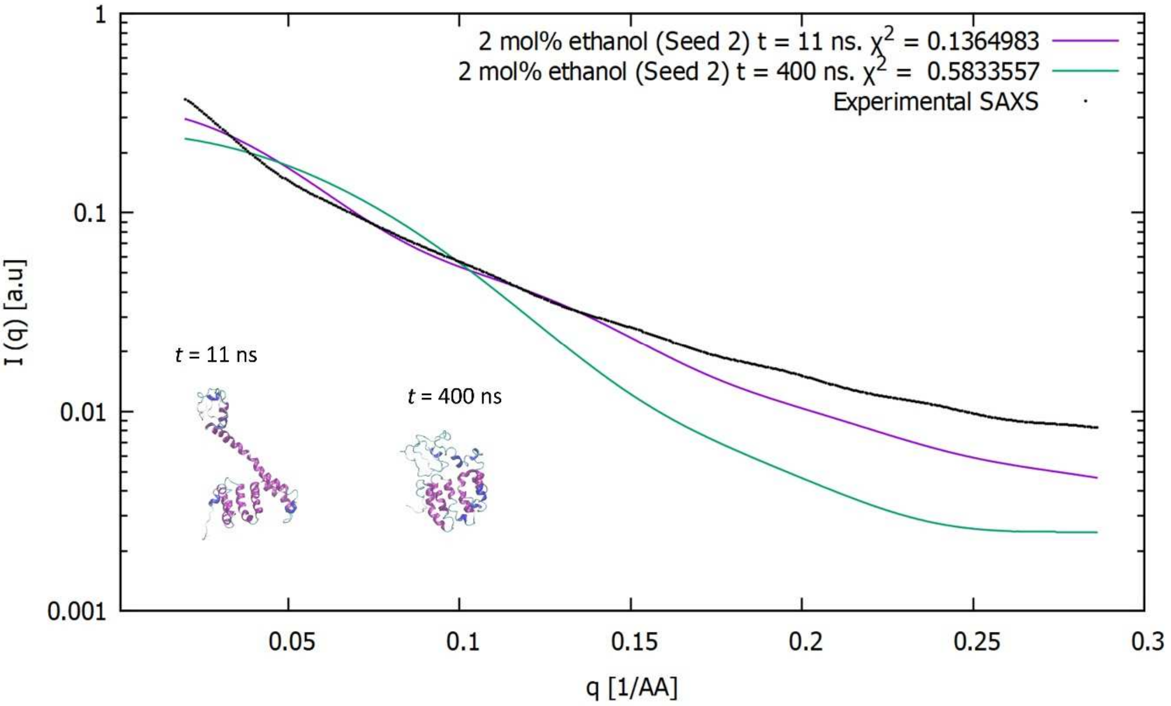


**SAXS predictions.** Predictions for 2 mol% ethanol showing the best fit (*t* = 2 ns) and the fit to the last MD frame (*t* = 400 ns).
