## Supplementary figures and images for "Conformations of a highly expressed Z19 α-zein studied with AlphaFold2 and MD simulations"

### S18 Fig

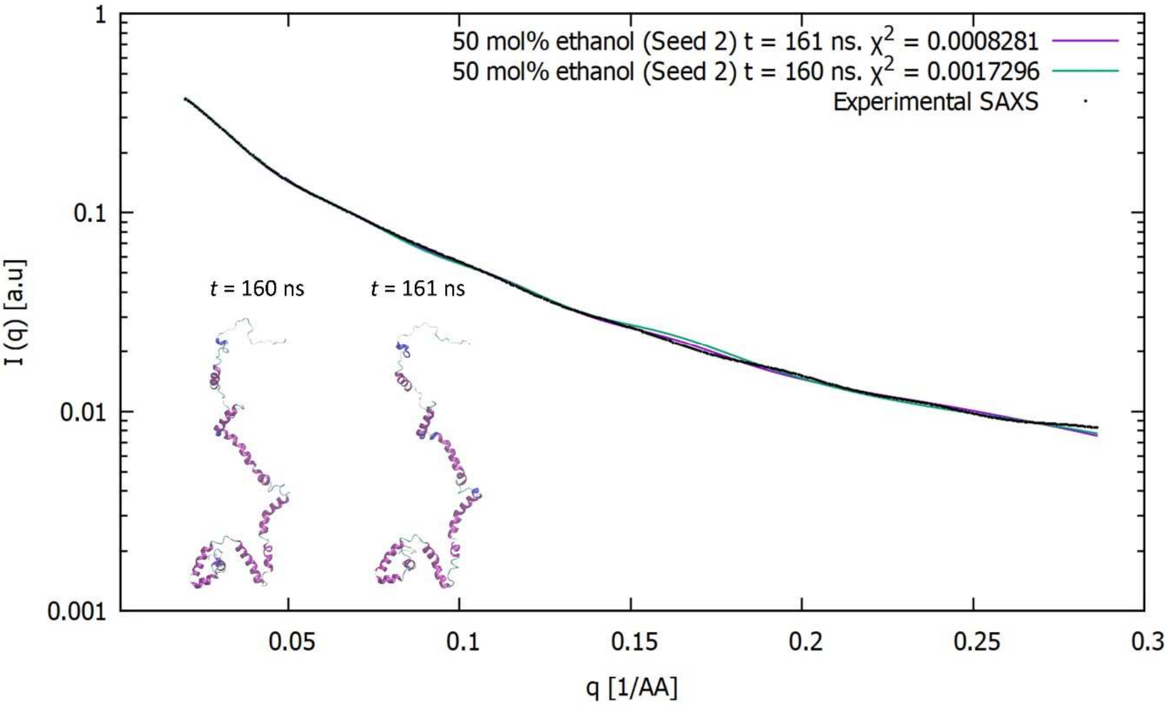


**SAXS predictions.** Predictions for 50 mol% ethanol showing the two best fits at $t$ = 161 ns and $t$ = 160 ns.

### S19 Fig

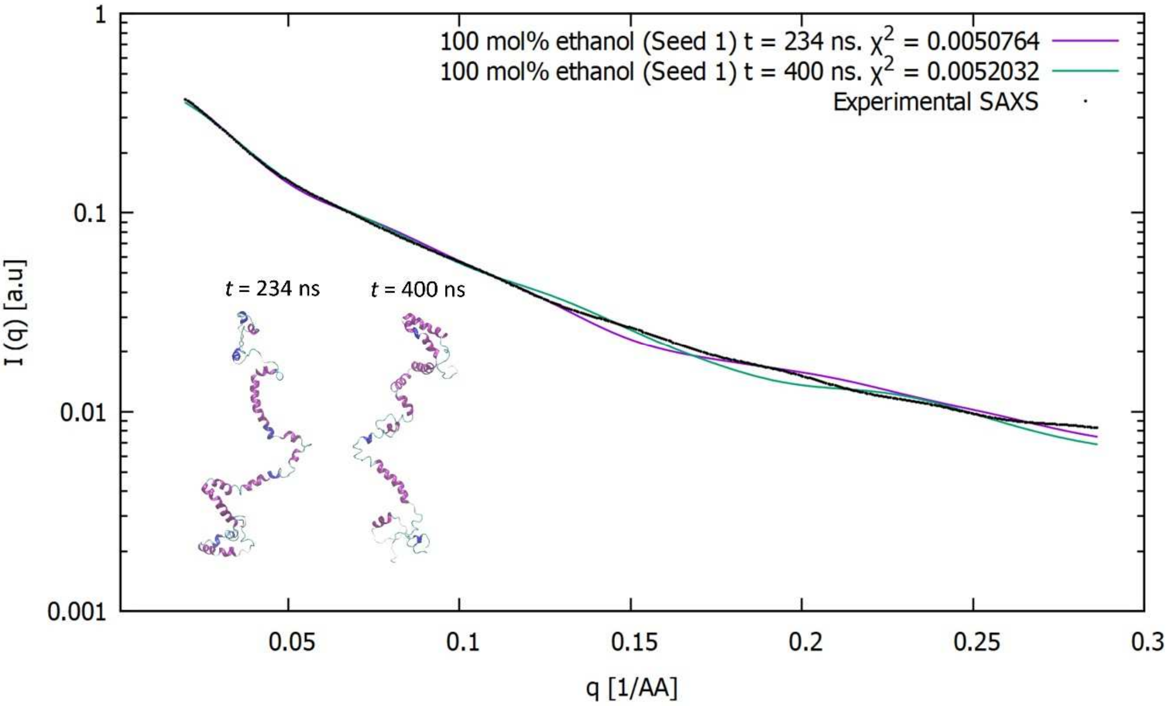


**SAXS predictions.** Predictions for 100 mol% ethanol showing the two best fits at *t* = 234 ns and *t* = 400 ns.

### S35 Fig

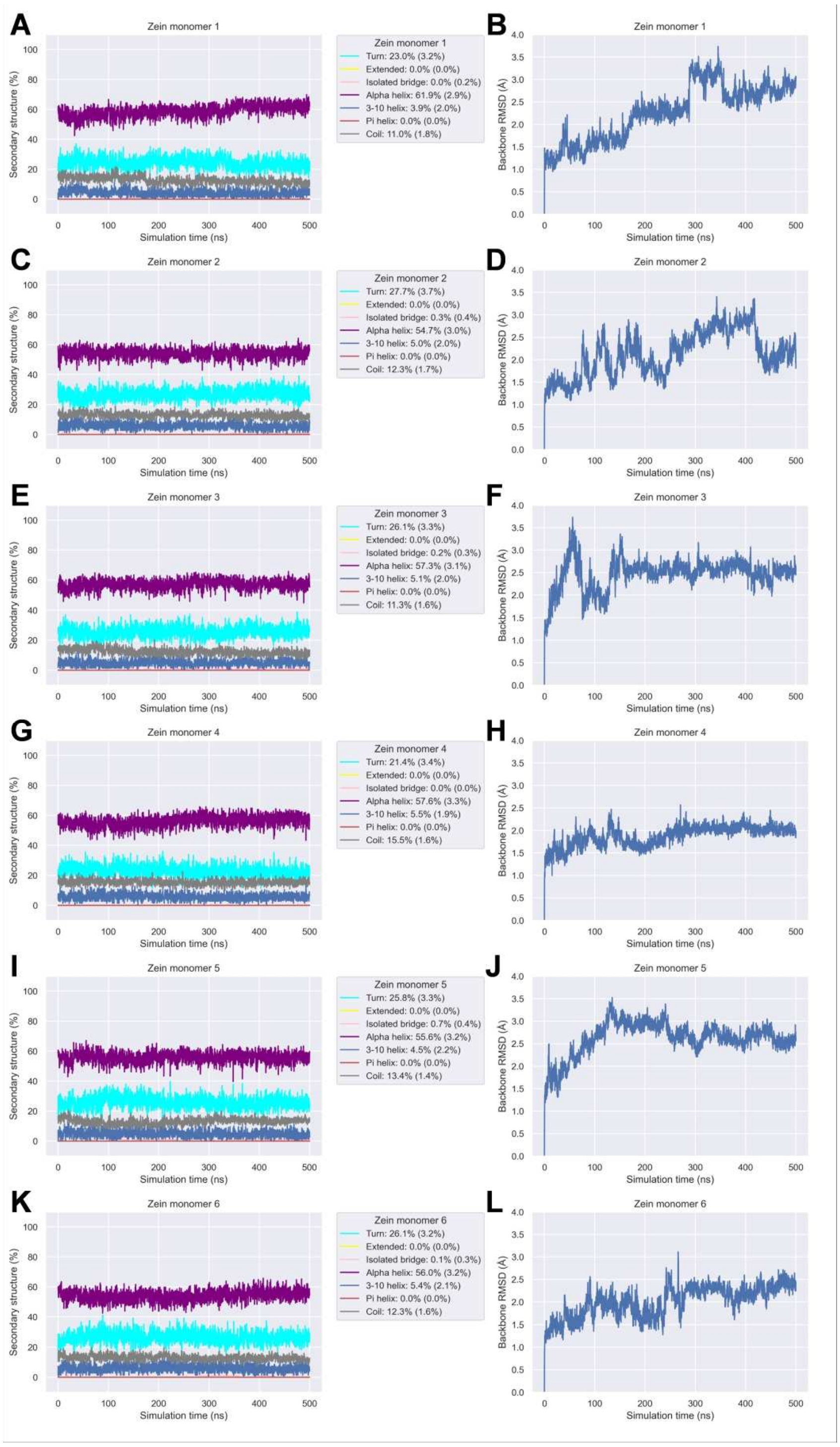


**Secondary structure and backbone RMSD for all-atom MD simulations of 6 zein copies in a water box.**
