## Supplementary material for "Conformations of a highly expressed Z19 α-zein studied with AlphaFold2 and MD simulations": S21 Fig

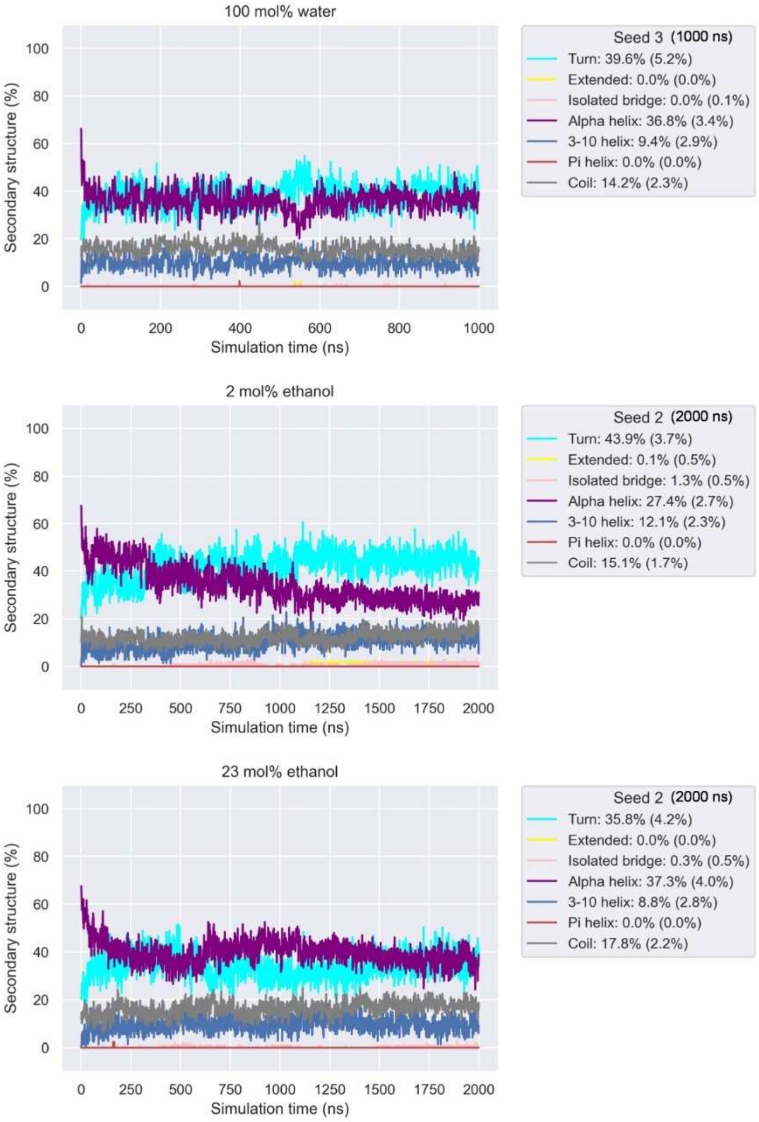


**Time series for secondary structure for extended GROMACS all-atom MD simulations.** Top: 100 mol% water simulation extended to 1000 ns. Middle: 2 mol% ethanol simulation extended to 2000 ns. Bottom: 23 mol% ethanol simulation extended to 2000 ns. Values given in the legend are averages over the last 100 ns of each simulation, with standard deviations given in brackets.
