## Supplementary material for "Conformations of a highly expressed Z19 α-zein studied with AlphaFold2 and MD simulations": S23 Fig

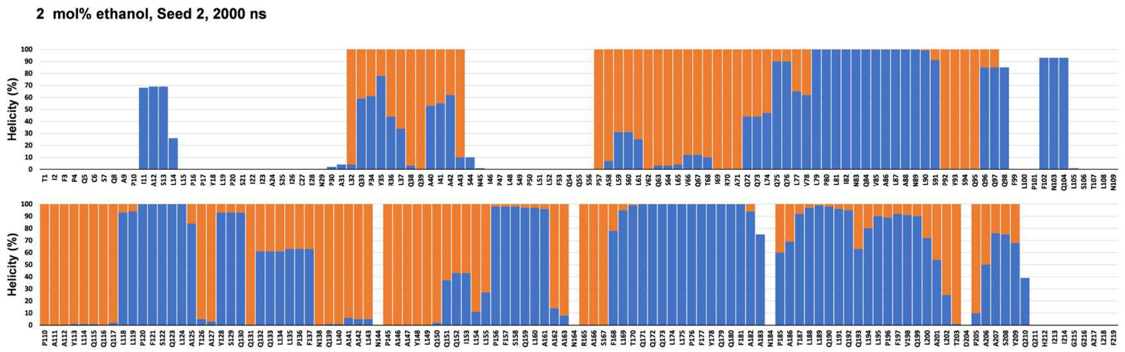


**Helicity per residue (sum of $3_{10}$- and α-helicity assigned by STRIDE) for the 2 mol% ethanol 2 μs simulation.** Orange: Helicity in the initial AlphaFold2 model. Blue: Helicity during the last 100 ns.
