## Supplementary material for "Conformations of a highly expressed Z19 α-zein studied with AlphaFold2 and MD simulations": S25 Fig

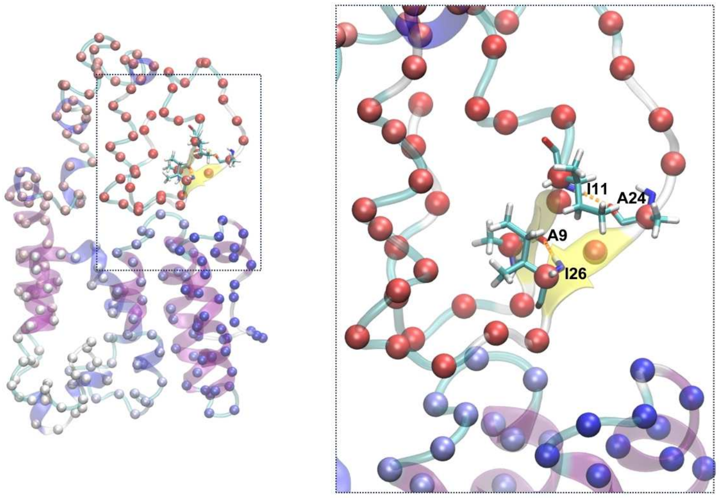


**Example of β-sheet structure in the extended 2 mol% (seed 2) ethanol simulation.** Left: MD snapshot at 1971 ns. Right: Enlarged view of the backbone hydrogen bond pairs A9/I26 and I11/A24. The protein backbone is shown in ribbon representation coloured by secondary structure. α-carbons are shown as spheres coloured by sequence position (red: N-terminal, blue: C-terminal).
