## Supplementary material for "Conformations of a highly expressed Z19 α-zein studied with AlphaFold2 and MD simulations": S26 Fig

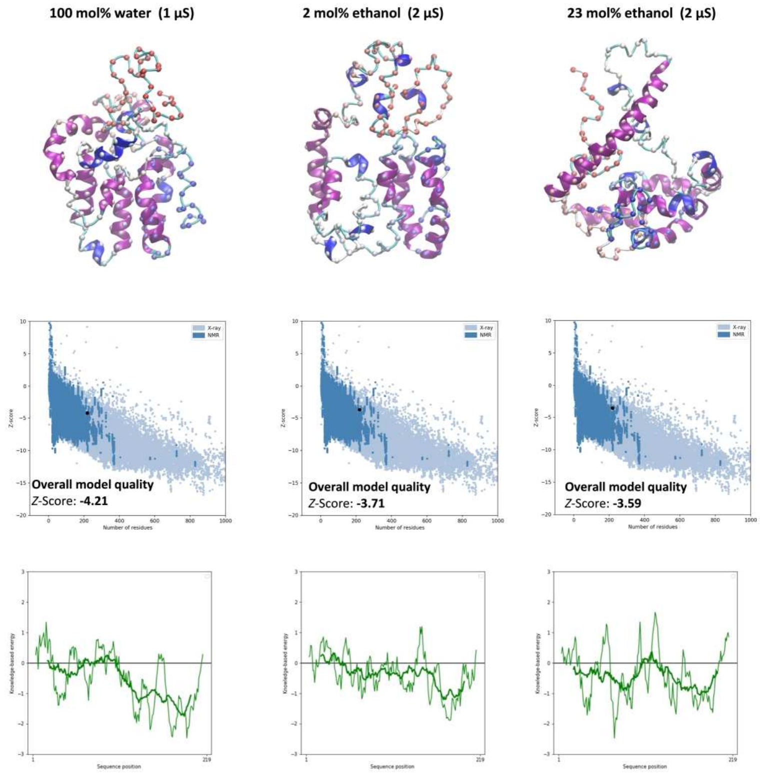


ProSA-web [1] quality checks for extended MD simulation endpoints.

### References


1. Wiederstein M, Sippl MJ. ProSA-web: interactive web service for the recognition of errors in three-dimensional structures of proteins. Nucleic Acids Research. 2007;35(suppl_2):W407-W10. doi: 10.1093/nar/gkm290.
