## Supplementary material for "Conformations of a highly expressed Z19 α-zein studied with AlphaFold2 and MD simulations": S27 Fig

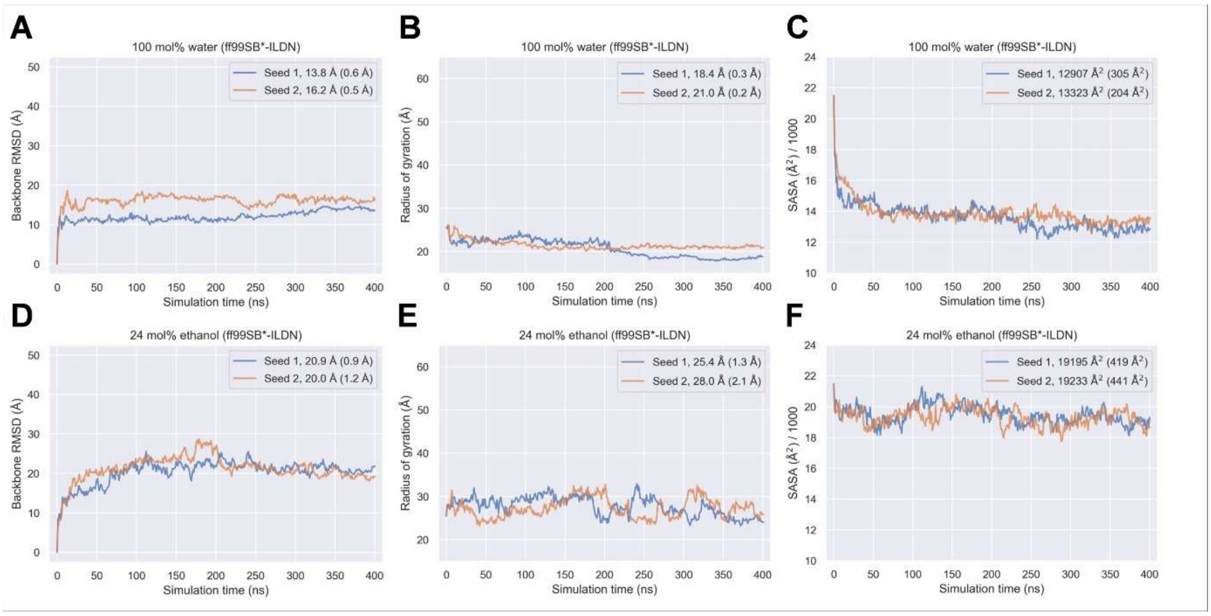


**MD time series for 400 ns GROMACS all-atom MD simulations with the ff99SB*-ILDN force field.** A, D: Backbone RMSD. B, E: Radius of gyration. C, F: SASA. Values given in the legend for each seed are averages over the last 100 ns of each simulation, with standard deviations given in brackets.
