## Supplementary material for "Conformations of a highly expressed Z19 α-zein studied with AlphaFold2 and MD simulations": S28 Fig

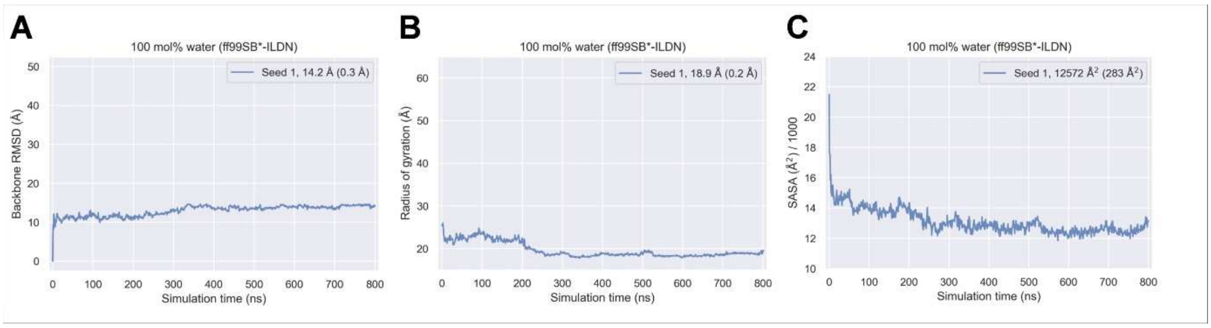


**MD time series for an extended GROMACS all-atom MD simulation with the ff99SB*-ILDN force field.** A: Backbone RMSD. B: Radius of gyration. C: SASA. The value given in the legend is an average over the last 100 ns, with standard deviation given in brackets.
