## Supplementary material for "Conformations of a highly expressed Z19 α-zein studied with AlphaFold2 and MD simulations": S29 Fig

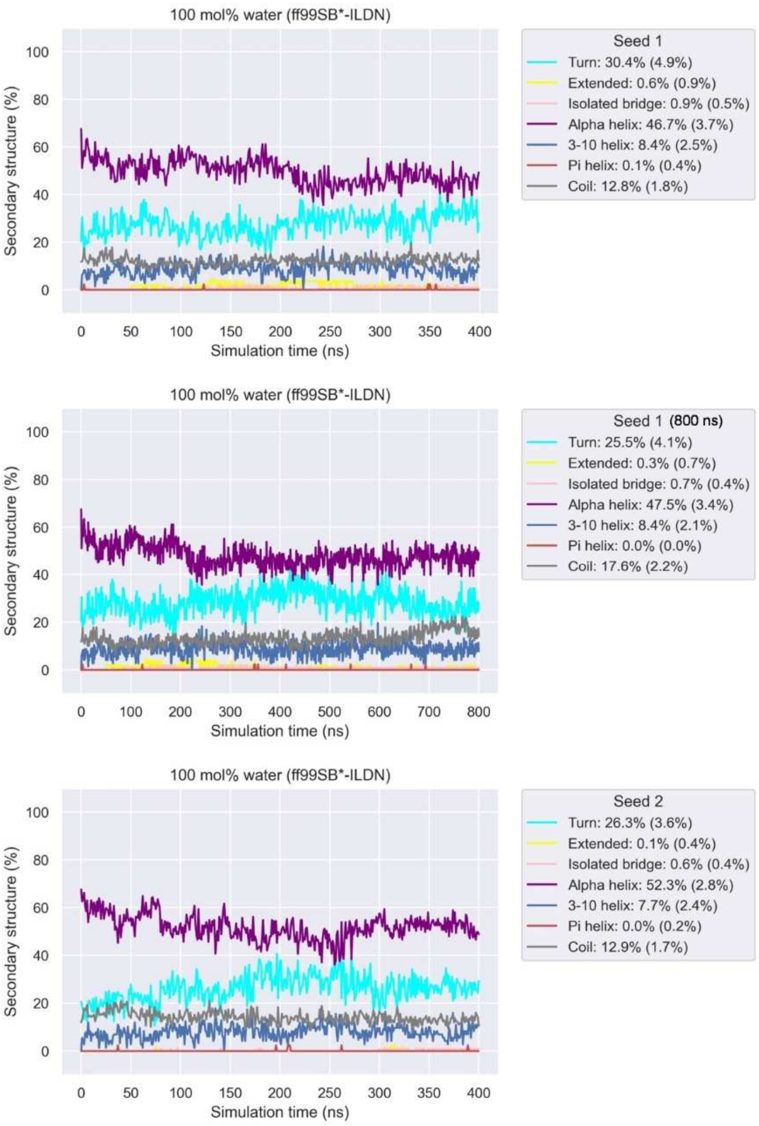


**Time series for secondary structure for 400 ns GROMACS all-atom MD simulations with the ff99SB*-ILDN force field.** Top: 100 mol% water simulation (seed 1). Middle: 100 mol% water simulation (seed 1) extended to 800 ns. Bottom: 100 mol% water simulation (seed 2). Values given in the legend are averages over the last 100 ns of each simulation, with standard deviations given in brackets.
