## Supplementary material for "Conformations of a highly expressed Z19 α-zein studied with AlphaFold2 and MD simulations": S30 Fig

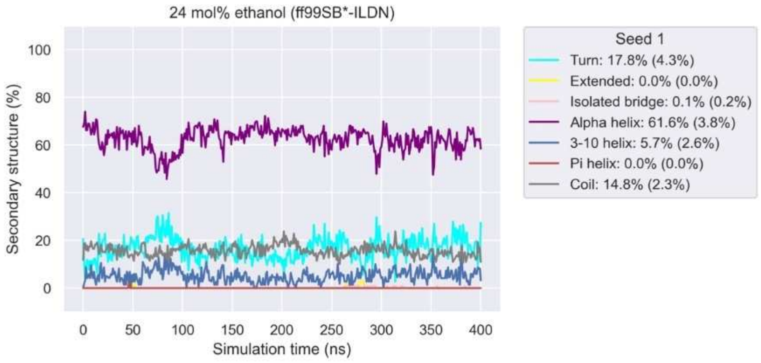


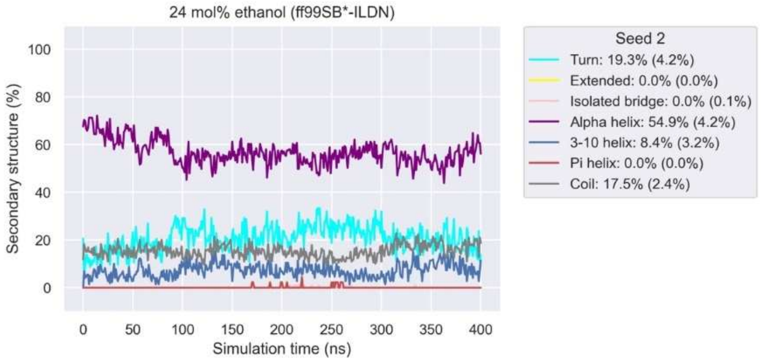


**Time series for secondary structure for 400 ns GROMACS all-atom MD simulations with the ff99SB*-ILDN force field.** Top: 24 mol% ethanol simulation (seed 1). Bottom: 24 mol% ethanol simulation (seed 2). Values given in the legend are averages over the last 100 ns of each simulation, with standard deviations given in brackets.
