## Supplementary material for "Conformations of a highly expressed Z19 α-zein studied with AlphaFold2 and MD simulations": S32 Fig

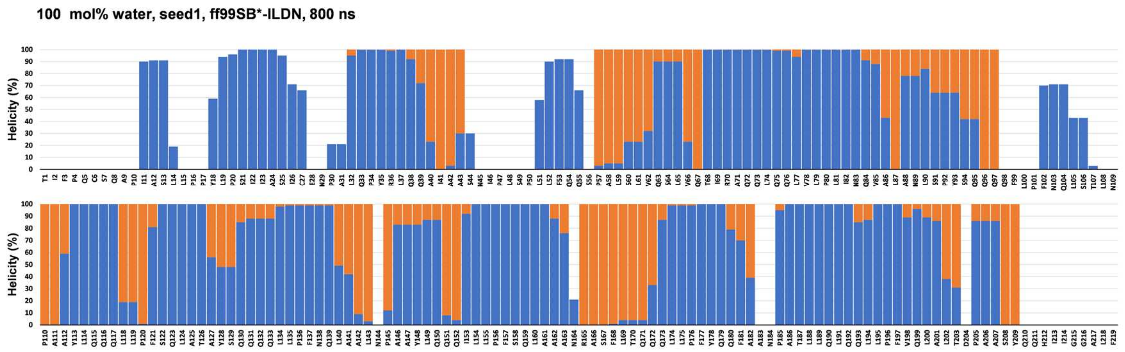


**Helicity per residue (sum of $3_{10}$- and α-helicity assigned by STRIDE) in the initial AlphaFold2 model and averaged over the last 100 ns of the 100 mol% water 800 ns MD simulation with the ff99SB*-ILDN force field.** Orange: Helicity in the initial AlphaFold2 model. Blue: Helicity averaged over the last 100 ns of the 100 mol% water Seed 1 MD simulation extended to 800 ns.
