## Supplementary material for "Conformations of a highly expressed Z19 α-zein studied with AlphaFold2 and MD simulations": S34 Fig

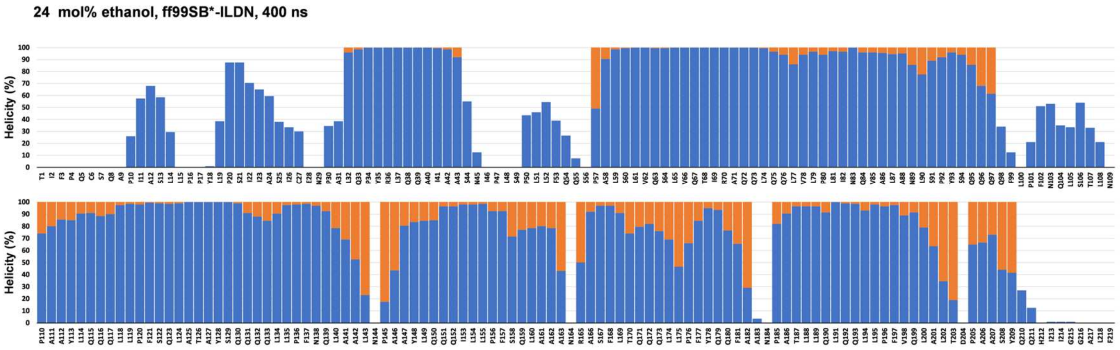


**Helicity per residue (sum of $3_{10}$- and α-helicity assigned by STRIDE) in the initial AlphaFold2 model and averaged over 24 mol% ethanol 400 ns MD simulations with the ff99SB*-ILDN force field.** Orange: Helicity in the initial AlphaFold2 model. Blue: Helicity of the last 100 ns averaged over the two (Seed 1, Seed 2) 24 mol% ethanol 400 ns MD simulations.
