## Supplementary material for "Conformations of a highly expressed Z19 α-zein studied with AlphaFold2 and MD simulations": S36 Fig

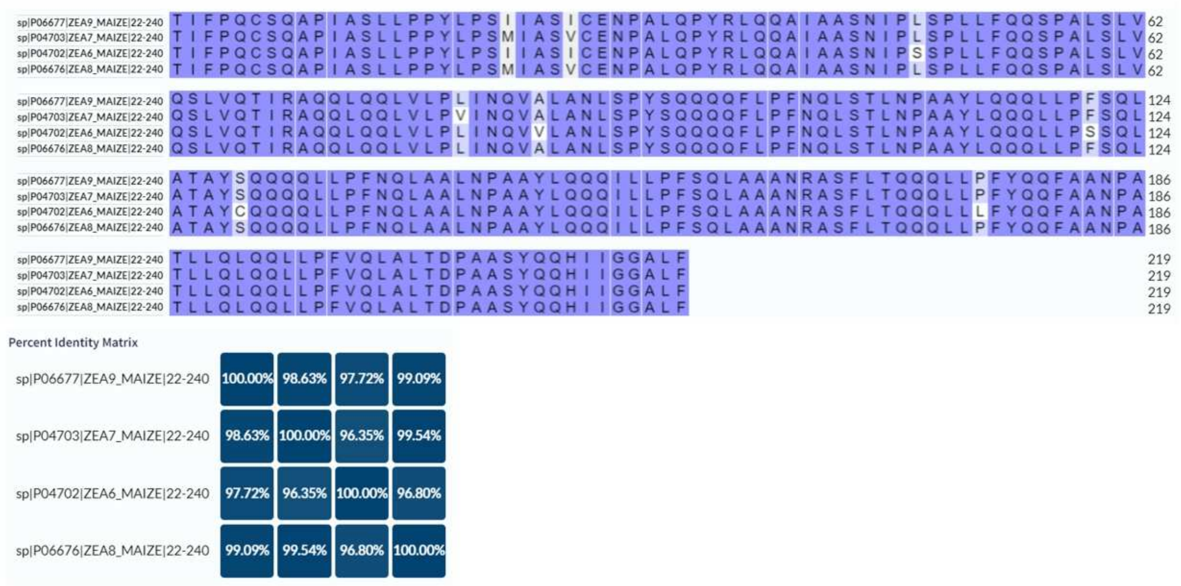


Mature protein sequence alignment and identity matrix for P06677 (19 kDa zein 19C2), P04703 (19 kDa zein A20), P04702 (19 kDa zein M6), P06676 (19 kDa zein 19C1).
